# Searching for universal model of amyloid signaling motifs using probabilistic context-free grammars

**DOI:** 10.1101/2021.01.22.426858

**Authors:** Witold Dyrka, Marlena Gąsior-Głogowska, Monika Szefczyk

## Abstract

**Background:** Amyloid signaling motifs are a class of protein motifs which share basic structural and functional features despite lack of apparent sequence homology. They are hard to detect in large sequence databases either with the alignment-based profile methods (due to short length and diversity) or with generic amyloid- and prion-finding tools (due to insufficient discriminative power). We propose to address the challenge with a machine learning grammatical model capable of generalizing over diverse collections of unaligned yet related motifs.

**Results:** First, we introduce and test improvements to our probabilistic context-free grammar framework for protein sequences that allow for inferring more sophisticated models achieving high sensitivity at low false positive rates. Then, we infer universal grammars for a collection of recently identified bacterial amyloid signaling motifs and demonstrate that the method is capable of generalizing by successfully searching for related motifs in fungi. The results are compared to available alternative methods. Finally, we conduct spectroscopy analyses of selected peptides to verify their structural and functional relationship.

**Conclusions:** While the profile HMMs remain the method of choice for modeling homologous sets of sequences, PCFGs seem more suitable for building meta-family descriptors and extrapolating beyond the seed sample.

## Background

Proteins forming amyloid structures have long been a subject of intensive research because of their association with the neurodegenerative diseases. Recently, there has been ever increasing interest in functional amyloids involved in normal physiological processes, for example, in establishing biofilms and membrane-less organelles, and in transmitting molecular signals. In general, the amyloids are defined in terms of the physical structure of cross-*β* polymer [1, 2, 3]. The essential feature of known amyloid proteins are short amino-acid motifs facilitating aggregation into a beta-sheet-like structure [4, 5]. The templating mechanism of forming amyloid fibrils can be exploited for acting as a prion. The prions are defined in terms of the function of infectious propagation. Indeed, amyloid proteins can act as prions capable of infectious propagation through imposing their own spatial structure on other proteins [3]. A well-known example is the [Het-s] prion from *Podospora anserina* [6, 7]. It is closely related to an ancient signaling pathway of which the amyloid-forming motif is a key element [8, 9, 3]: related motifs were identified in metazoa [10, 11], fungi [12] and bacteria [13]. At least some of these sequence motifs of roughly 20 amino acids form a beta arch fold [14] and often they contain polar amino acids: asparagine and glutamate [15]. Despite these common features, the already identified amyloid signaling motifs (ASM) in bacteria and fungi exhibit high sequence diversity beyond noticeable homology [12, 13]. Is it therefore possible to define universal rules to be obeyed by sequences of functionally-related yet non-homologous ASM? Such a model would allow to identify new amyloid signaling motifs in ever growing data sets of genomic sequences. Moreover, it could facilitate better understanding of mechanisms of conformation transmission and aggregation.

Evolutionary related sequence families are typically modeled with the profile Hidden Markov Models (pHMM) [16, 17]. As they assume sequence homology and rely their training process on the multiple sequence alignment (MSA), pHMMs are not best suited for modeling diverse collections and meta-family of motifs. Moreover, their discriminative power is limited, especially for short sequences, as amino acid distribution at each position in the alignment is modeled separately.

There exist also various methods dedicated to recognition of amyloidogenic regions of protein sequences [18, 19, 20, 21]. They are mainly based on statistical properties of hundreds of hexapeptides confirmed experimentally to aggregate into amyloid-like fibrils [22]. Unfortunately, these methods often fail to detect functional prion-related amyloid motifs. Indeed, it seems that the amino acid composition of prions has to differ from that of typical amyloids in order to balance water soluble and aggregated state in physiological conditions. This led to developing dedicated prion predictors including composition based PAPA [23] and more sophisticated pWaltz [15]. The latter is based on the model of prion sequence as an amyloidogenic core (or aggregation seed) within a disordered region. Yet, these methods still miss a considerable fraction of HET-s related ASM. One apparent feature of the motifs that is missing from these models is the propensity to forming the beta-arch structure. This is specifically addressed by ArchCandy [24], a method for detecting beta-arches in protein sequences.

In this piece of research we propose a method aiming at exploiting benefits of beta-arch detection and amyloidogenic composition in a single elegant model. We present a model based on the Probabilistic Context-Free Grammar (PCFG), which extends the profile Hidden Markov Model with capability to capture some dependencies between distant positions in the sequence [25, 26, 27, 28, 29, 30]. PCFG is well suited to model nested dependencies resulting from interactions between strands involved in the beta-turn-beta structures (cf. [31]), as recently demonstrated for the HET-s motif [32]. In the same work, the PCFG model was shown to be capable of generalizing between two apparently heterologous architectures of Calcium binding sites [32].

The PCFG model does not assume evolutionary relationship between the sequences, as it does not rely on the multiple sequence alignment. In [32], the grammars were trained with the genetic algorithm within a setup that significantly limited the number of rules and thus complexity of the model. Here, we propose to use statistical learning, the Inside-Outside (IO) algorithm [33] that allows for training much larger grammars. While the IO algorithm is considered more prone to converge to local minima [34], the benefit of extending the rule set many fold seems to be an overwhelming advantage when describing more complex languages.

The main contributions of the paper are as follows. First, we show that in grammatical modeling of protein sequences, statistical learning leads to comparable or better results than previously used evolutionary learning, while being incomparably quicker. Second, we report on a benefit of smoothing learned profiles of amino-acid emissions represented with the lexical rules. Third, we show that the PCFG model is capable of representing individual families of amyloid signaling motifs, and is practical in searching them in sequence databases. Fourth, we present our main result: the model that generalizes over various amyloid signaling motif families, obtained by training a common PCFG for a set of ten motif families from bacteria. This universal model is then validated by searching for already known fungal ASMs. Fifth, we experimentally verify spectroscopic characteristics of selected diverse motifs detected with the PCFG model.

## Methods

### Computational methods

Probabilistic Context-Free Grammar (PCFG) is a generative probabilistic model of sequential categorical data [25]. Under the model, sequences are derived from the start symbol using rewriting rules, associated with some probabilities, until all remaining symbols are non-derivable (or terminal). Formally, PCFG is a quintuple 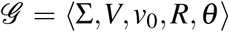, where Σ (alphabet) is a set of terminal symbols, *V* is a set of non-terminal symbols (variables) disjoint from Σ, *v*_0_ ∈ *V* is a start symbol, *R* is a set of production rules rewriting variables into strings of variables and/or terminals, and *θ* is a set of corresponding rule probabilities. An illustrative toy example of PCFG modeling a subfamily of beta-hairpin protein sequences can be found in [32]. A probabilistic grammar is *proper* if rule probabilities sum up to 1 over rules rewriting the same variable. A complete derivation is a chain of rules beginning with *v*_0_ and finishing with a string of terminal symbols. Each derivation can be represented as a parse tree. The probability of derivation is the product of probabilities of rules involved. In turn, probability of a sentence *x* given 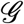 is the sum over all derivations that generate *x*. The grammar is called *consistent* if the probability mass distributed by the grammar over all sentences sums up to 1. Language is a set of all sentences that can be derived according to the grammar.

Each context-free grammar (whether probabilistic or not) that does not generate empty sentences can be translated to the Chomsky Normal Form (CNF) [35]. This canonical form implies that production rules are either in the form *A → a* (lexical rules) or *B → CD* (structural rules), where lowercase letters denote terminal symbols, while uppercase — non-terminal symbols. In addition, each CNF grammar that does not generate single letter sentences can be translated to the form where the sets of variables that can be rewritten with the lexical and the structural rules are disjoint. We call such a grammar form *bipartite* CNF, and denote the two groups of variables as lexical and structural non-terminals, respectively [36].

Context-free grammars are suitable to represent branching and nesting in syntactic description of sequences, but CNF makes the latter unnecessarily lengthy. It is therefore convenient to extend the CNF with *contact* rules, which rewrite variables into triples made of lexical, structural, and lexical non-terminals. This grammar form called Chomsky Form with Contacts (CFC) is especially suitable to represent pairs of amino acids in contact [32]. Since spatial proximity of residues often generates mutual constraints, using a contact rules that generates both residues at once is an effective way to model their dependency.

A parse tree generated with CFG for a protein sequence can be compared to the contact map if corresponding spatial structure is available [36]. However, the PCFG sums the probabilities over all parse trees derivable for the given sequence. Indeed, the most likely parse tree often does not approximate well the most likely shape of all such parse trees [37]. Fortunately, in the case of grammars in the CFC form, it is possible to calculate the probability map of parsing a pair of residues through the contact rules (referred later as probability map of pairing). It can be reasonably expected that residues in contact are often generated with the contact rules. Instead of searching for the most likely shape of parse trees, as used in the RNA structure prediction [38], we propose to compare the probability map of pairing for the best matching sequence fragment with its corresponding spatial distance map.

Probabilities of PCFG rules can be inferred from a positive training set of sequences using the Inside-Outside (IO) algorithm [33], which implements the Expectation-Maximization scheme [39]. The algorithm can quickly handle thousands of rules but is prone to converge to local minima [40]. When applied to large generic set of rules constituting a *covering grammar*, the process of optimizing rule probabilities of which most eventually become zero, is akin to learning grammar. The most popular alternatives to IO are based on Genetic Algorithms (GA) using either a fixed set of rules [41, 29], as in the case of IO, or learnable set of rules [42, 43, 34, 44].

Efficiency of learning PCFG can be improved when syntactic trees [26, 27, 45, 46, 47] or partial syntactic constraints are available [48, 49]. Recently, we proposed using pairwise contacts between amino acids to constrain the GA-based learning of PCFG in CFC form for protein motifs. We showed that even a few relevant contacts led to learning better performing grammars [32].

The outline of the processing pipeline, as proposed and tested in this study, is provided in Supplementary Fig. 1. Newly added features are described in the Results section.

### Materials

Computational experiments were carried out using several sets of protein sequences. The collections included existing samples, which were used to benchmark the improved method against the previous approach, and novel samples of diverse amyloid signaling motifs, which were used to test capability of the current method to generalize.

#### CaMn

A benchmark set of 24 sequences of a Calcium and Manganese binding site from the legume lectins [50] was collected according to PROSITE pattern PS00307 [51] true positive and false negative hits, extended to 27 residues to cover the entire binding site, as in [36, 32]. The motif folds into a stem-like (beta-loop-beta) structure with over 40 internal contacts, many of them forming nested dependencies keeping together beta-strands at the ends of the motif [52]. The sequences were made non-redundant at identity of 70% (nr70) using cd-hit [53].

#### HET-s

A benchmark set of 160 sequences (nr70) of the HET-s-related motifs r1 and r2 involved in the prion-like signal transduction in fungi was derived from [54]. The largest subset of motifs with length of 21 amino acids was used, as in [36, 32]. The beta-hairpin-like fold of the motif partially relies on interactions between hydrophobic amino acids. HET-s motifs r1 and r2 are known to adopt the beta-hairpin-like fold when templated by the related motif r0 located in the N-terminus of a cooperating NLR protein [55]. While the r0 motifs share a considerable sequence similarity with the interacting r1 and r2 motifs (average identity of around 30%), they contain significantly less aspartic acid, glutamic acid and lysine, and more histidine and serine [54]. A set of 98 HET-s r0 motifs was manually extracted from genes of NLR proteins adjacent to genes encoding proteins containing the r1 and r2 motifs [54]. To test sensitivity of the models trained for the r1 and r2 motifs, we used a subset of 77 non-redundant 21-residue long r0 motifs, as in [32]. HET-s is the only analyzed motif with experimentally solved structure. A high-resolution NMR structure of HET-s amyloid fibrils made of the effector-side r1 and r2 motifs in *Podospora anserina* sequence Q03689 is available in the Protein Data Bank (pdb: 2kj3) [56].

#### BASS

Novel families of bacterial amyloid signaling motifs, termed BASS 1 to 10, were identified in neighboring C-termini of Bell domain homologs and N-termini of NLR proteins in bacteria [13]. Each family was defined according to a set of related profile HMMs. For the current piece of research, for each motif family we extracted fragments of Bell-side sequences matched by the motif profile HMMs. Then, we aligned them using Clustal Omega [57] with the *–auto* parameter and submitted to Gremlin [58] to obtain contact constraints. Residue-residue contacts were found for all but three families with the least effective number of sequences (BASS 7, 8, 10). Considering only the most reliable contact pairs, we hand-crafted contact constraints including from 1 to 5 non-overlapping (hence context-free compatible) pairs of residues in contact. We then mapped the contacts onto unaligned sequences. For training, we used the Bell-side motifs samples (nr70) including from 329 (BASS2) to only 7 (BASS10) sequences with length varying from 20 to 40 amino acids. For testing, we used the N-termini of NLR proteins with 297 known instances of motifs BASS1-10, according to Supplementary Table 2 in [13].

#### Other BASS

In our previous research [13], a number of pairs of similar amyloid-like patterns were identified in C- and N-termini of neighboring Bell and NLR proteins while being missed with profile HMMs (see Supplementary Table 2 in [13]). We extracted 100 amino-acid long Bell C-termini and 150 amino-acid long NLR N-termini containing these *other* BASS motifs, and made them non-redundant (nr70). This yielded a set of 18 Bell-side C-termini and 26 NLR-side N-termini, which was used for further testing. Interestingly, the *BASSother* set included 3 sequences from the Archaea species.

#### Fungal test motifs

Test sets were made of fungal amyloid signaling motifs [12] extracted from a recent set of NLR proteins [59]. The sets included sequence fragments matching Pfam profiles of motifs sigma (Pfam NACHT_sigma, 20 sequences), HET-S (Pfam HET-S, 12 sequences) and PP (Pfam Ses_B, 22 sequences), which were made non-redundant at identity of 70%.

#### PDBfull and PDBfrag

The first negative sample was designed to rather roughly approximate the entire space of protein sequences. It was based on the negative set from [29], which consisted of 829 single chain sequences of 300-500 residues retrieved from the Protein Data Bank [60] at the identity threshold of 30% (accessed on 12th December 2006). In addition, we used the negative sample obtained by cutting the basic negative set into overlapping subsequences of the maximum length of positive sequences and made non-redundant at identity of 70%, as in [32].

#### NLReff

The second negative set was based on a sample of 7901 NLR proteins with N-terminal known to contain non-prion-forming effector domains [59] except for the PNP_UDP_1 domain (see below). The actual negative set consisted of over 2411 fragments matching the Pfam profiles of the domains and non-redundant at identity of 70%. Length of the fragments ranged from 41 to 366 amino acids (median: 175). The set was designed to approximate the background encountered when finding positive test motifs in their typical setting in the N-termini of NLRs. The restriction to include only boundaries of domain profiles was based on the fact that putative functional amyloid motifs are sometimes present between the effector and nucleotide-binding domain.

PNP_UDP_1 domains were excluded from the set because of a part of their sequences predicted to be amyloidogenic according to PASTA2 and AmyloGram, and involved in the beta arch according to ArchCandy. Indeed, available structures show that the fragment consists of two beta strands connected with a loop (e.g. positions 60–100 in pdb:1zos) [61].

#### DisProt

The third negative set was adopted from evaluation of the ArchCandy tool [24] and consisted of 48 sequences (nr70) of soluble disordered protein regions without link to amyloidoses. The set originated from the DisProt database [62]. Length of fragments ranged from 37 to 149 amino acids (median: 101.5). The set was used to check specificity of the tested models against non-amyloidogenic disordered proteins.

#### Peptides selected for experimental verification

Out of sequence fragments identified as ASMs with grammars and other existing computational methods, we selected four peptides for experimental verification if they form structures consistent with expected features (presence of the beta arch and amyloid-like aggregation). First, to twick and test the experimental setup, we used two *bona fide* effector-side ASM peptides: BASS3 RHIM-like motif from *Frankia sp.* ORT49035.1 (positions 103 to 123) [63, 13] and *Nectria haematococca* sigma motif from AAS80314.1 (349 to 385) [64, 12]. Then we analyzed *Methanothrix soehngenii* AEB69175.1 (5 to 29) [65]. This archeal NLR-side motif resembling BASS3 was originally identified through local pairwise alignment of proteins coded by neighboring genes while being missed with the profile HMM-based search [13]. Finally, we experimentally verified a sequence fragment resembling the sigma motif in *Coleophoma crateriformis* NLR protein RDW70414.1 (382 to 421) [66].

### Experimental methods

#### Peptide synthesis

All commercially available reagents and solvents were purchased from Lipopharm.pl, Sigma-Aldrich and Merck and used without further purification. Peptides were obtained with an automated solid-phase peptide synthesizer (Liberty Blue, CEM) using rink amide AM resin (loading: 0.59 mmol/g). Fmoc deprotection was achieved using 20% piperidine in DMF for 1 min at 90 °C. A double-coupling procedure was performed with 0.5 M solution of DIC and 0.25 M solution of OXYMA (1:1) in DMF for 4 min at 90 °C. Cleavage of the peptides from the resin was accomplished with the mixture of TFA/TIS/H_2_O (95:2.5:2.5) after 3 h of shaking. The crude peptide was precipitated with ice-cold Et_2_O and centrifuged (8000 rpm, 15 min, 2 °C). Peptides were purified using preparative HPLC (Knauer Prep) with a C18 column (Thermo Scientific, Hypersil Gold 12 *μ*, 250 mm × 20 mm) with water/acetonitrile (0.05% TFA) eluent system.

Analytical high-performance liquid chromatography (HPLC) was performed using Kinetex 5*μ* EVO C18 100A 150 × 4.6 mm column. Program (eluent A: 0.05% TFA in H_2_O, eluent B: 0.05% TFA in acetonitrile, flow 0.5 mL/min): A: t=0 min, 90% A; t=25 min, 10% A.

Peptides were studied with WATERS LCT Premier XE System consisting of high resolution mass spectrometer (MS) with a time of flight (TOF).

#### Amyloid-like aggregation

To determine aggregation properties of studied peptides Attenuated Total Reflection – Fourier Transform Infrared (ATR-FTIR) experiments were carried out. Vibrational spectroscopy is widely used for protein and polypeptides secondary structure analysis [67, 68] and for monitoring the aggregation processes in amyloids studies [69, 70, 71]. The Amide I band (1700–1600 cm^−1^) corresponding to C=O stretching vibrations and the Amide II band (1600-1500 cm^−1^) arising mainly from in-plane N-H bending of the peptide bonds are the most useful for secondary structure estimation. For *α*-helical proteins the maxima of Amide I and Amide II bands are observed at around 1655 cm^−1^ and 1545 cm^−1^, respectively. Random structures possess the Amide I located at 1645 cm^−1^. Native *β*-sheet rich proteins show amide bands maxima near 1635 and 1530 cm^−1^. When the aggregation occurs, the Amide I band is narrowing and shifting to lower wavenumbers. Rigid and highly ordered amyloid fibrils exhibit the Amide I band below 1625 cm^−1^ [69]. The high water absorption in the Amide I region is main drawback of IR spectroscopy. Subtracting the water absorption spectra may cause significant distortion to the spectral line shape [72]. That is why deuterium oxide is used as an alternative solvent. Due to the frequency of the OH bending mode of D_2_O molecules is lowered (from 1635 cm^−1^ for H_2_O) to 1210 cm^−1^. Substitution of water by heavy water causes a relatively small down shift of Amide I band [67].

#### Spectroscopy

For spectroscopic measurements peptides were dissolved in D_2_O (deuterium oxide, 99,8% D, Carl Roth, GmbH, Germany) to a final concentration of 2 mg/mL. Peptide solutions were incubated at 37 °C (98.6 °F) for 24 h. ATR-FTIR spectra were collected using a Nicolet 6700 FT-IR Spectrometer (Thermo Scientific, USA) with Golden Gate Mk II ATR Accessory with Heated Diamond Top-plate (PIKE Technologies). The spectrometer was continuously purged with dry air. All spectra were obtained in the range of 4000−400 cm^−1^. Directly before sampling, the background spectrum of diamond/air was recorded as a reference (512 scans, 4 cm^−1^). Spectroscopic measurements were performed at air-dried peptide films. Initially, 10 *μ*l of peptide solution was dropped directly on the diamond surface and was allowed to dry out. For each spectrum 512 interferograms was coadded, with 4 cm^−1^ resolution. All spectra were registered at temperature of 37 °C (98.6 °F).

All spectra were analyzed using the OriginPro (version 2019, OriginLab Corporation, USA). The analysis included spectra baseline correction, smoothing using the Savitzky-Golay [73] polynomial filter (polynomial order 2, a window size of 31 points), normalization to 1 for the Amide I’ band and deconvolution into subcomponents using the Lorentz function based on second derivative spectra. ATR-FTIR spectra were initially preprocessed using OMNIC™ software (version 8, Thermo Fisher Scientific, USA): atmospheric and ATR correction.

## Results

In a previous paper [32], we showed that fungal prion-forming HET-s motifs r1 and r2 can be accurately represented with automatically inferred probabilistic context-free grammars comprising just of three lexical and four structural symbols (*l3s4*). In the cross-validation scheme, the model achieved the average precision (AP) of 0.60 for the negative to positive sample cardinality ratio over 200:1 [32]. Moreover, we found that consensus predictions from grammars comprising of seven structural symbols were practically useful for identifying related HET-s r0 motifs in NLR proteins with AP of 0.82 [32]. However, the PCFGs were outperformed in this task by less expressive profiles of Hidden Markov Models (pHMM). The presumed disadvantage of our PCFG approach was in likely over-simplicity of grammars due to the tiny number of non-terminal symbols. This limit was necessary to make the number of rules manageable in our GA-based scheme for inferring rule probabilities. Indeed, the number of possible rules increases exponentially with the number of available symbols and our implementation of evolutionary approach could not effectively handle search spaces of more than 500-1000 rules [32].

### Improved modeling of individual families of protein motifs

Even though the evolutionary scheme could be adjusted [74, 75], in the current project we resorted to the classical statistical learning method, the Inside-Outside (IO) algorithm [33]. The main advantage is relatively quick convergence of the procedure, even for hundred thousands of rules, thus allowing for considerably more non-terminals symbols (for example 40) in the *covering grammar* made of all possible rules.

#### Smoothing

The large size of such a grammar increases the risk of over-generalization, i.e. over-fitting rule probabilities to training data. A viable solution consists on smoothing the probabilities in the course of post-processing, so the grammar can parse sequences with amino acids unseen in the given context during training. While generally not trivial, the smoothing is relatively straightforward when applied to lexical rules modeling amino-acid emissions from lexical variables. Indeed, one can apply one of the classical mutation models such as PAM [76] or BLOSUM [77]. While the latter is typically considered more accurate, the former has the advantage of intrinsic simplicity and elegance of the underlying Markov model and thus was chosen for this project. In our implementation, distributions of amino acids modeled by lexical rules can be smoothed according to the requested number of point accepted mutations.

#### Cross-validation

The updated IO-trained PCFG method was tested on two benchmark sets from our previous research, the HET-s motifs r1/r2, and a Calcium and Manganese binding site motif (CaMn). Replicating the procedure from [32], we used a variant of the 8–fold Cross–Validation scheme in which 6 parts were used for training, 1 part was used for validation and parameter selection, and 1 part was used for final testing (the scheme resulted in 56 runs for each sample).

#### Learning efficiency of IO and GA

We first compared efficiency of evolutionary and statistical learning of grammars for the simpler CaMn set. Similarly to [32], we stuck to the *l3s4* grammars in the CFC form and compared training with and without contact constraints. Setups of evolutionary (GA) and statistical (IO) learning schemes were essentially identical except that the convergence criterion was calculated over 100 epochs for the former and just 10 iterations for the latter, simply because the Inside-Outside procedure converged much quicker: while on average 3000-4000 steps were needed using GA, only 10 or 40 iterations was enough for IO (Fig. 1a). Contact constraints seemed to shorten the training for IO but not for GA. When tested on the validation set against *PDBfrag*, best grammars achieved the maximum AP around 0.95 for evolutionary and 0.85 for statistical learning. This likely resulted from more pronounced over-fitting with IO, where the gap in performance between the training and validation sets was around 0.10-0.15, in comparison to 0.05 for GA (in terms of AP). Closer inspection of the induced best grammars revealed that the Inside-Outside was more prone to suppressing rule probabilities: around half of them was set below 1*e* − 5, and a couple of percent were set to zero. In contrast, the evolutionary scheme suppressed less than 1% below 1*e* − 5 and did not turn off any rule completely. (Of note is that in practice the structural rules with probability below 1*e* − 5 have at best negligible impact on the sequence probability log scores and can be pruned off in order to improve the speed of parsing.) This finding led us to trying the PAM-based smoothing on the lexical rules probabilities. This turned out to be highly efficient: the average precision of apparently over-fit IO-trained grammars was pushed to over 0.97, almost closing the performance gap of around 0.30 (for the longest training). The best results of smoothing were obtained with PAM values in the range of [5, 20] (Fig. 1b shows results for PAM10). For grammars trained with GA long enough to experience over-fitting, the smoothing pushed the average precision up to above 0.98.

**Figure 1:**
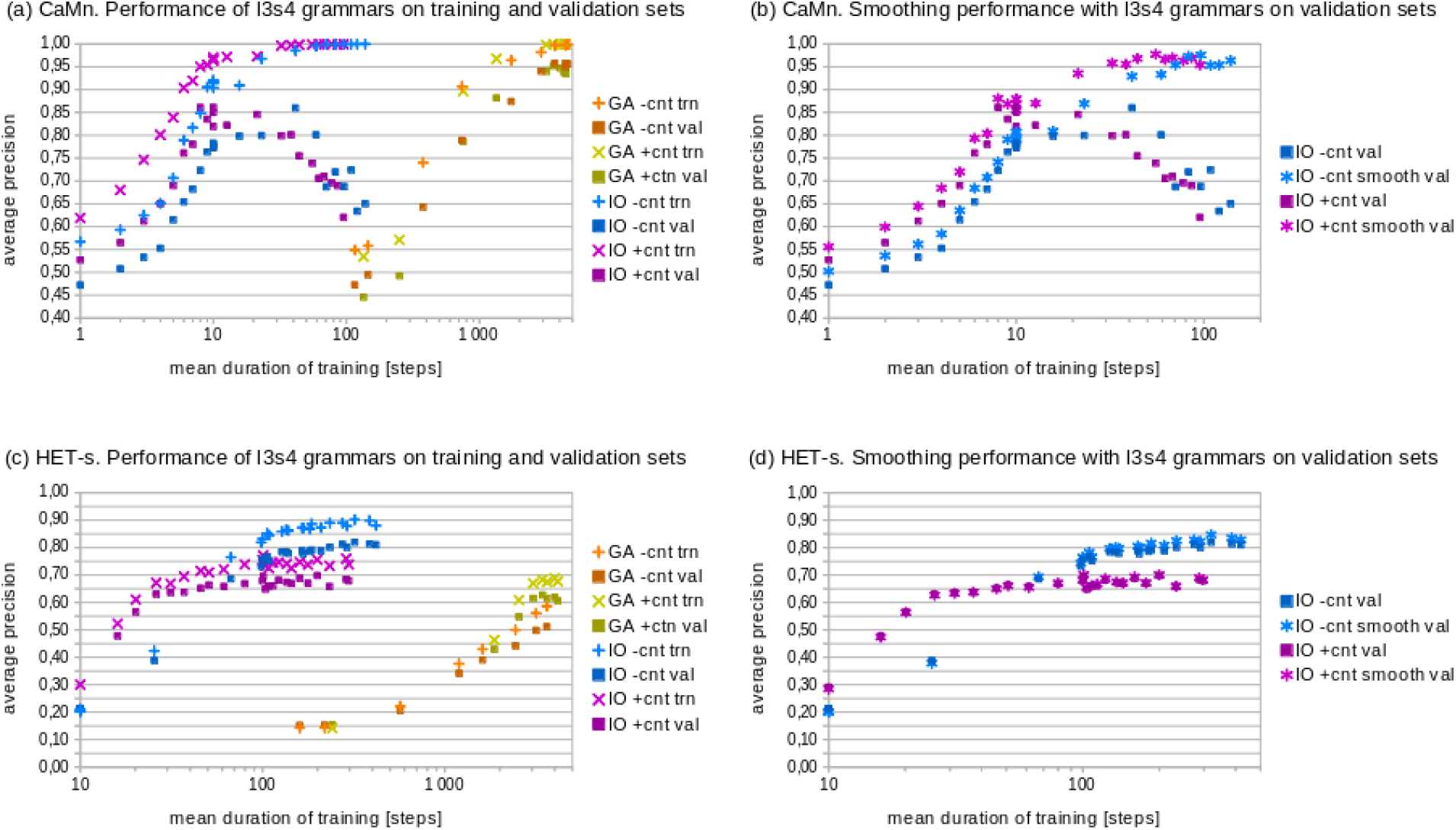
Performance comparison of grammar learning schemes. Mean average precision with regard to mean training duration achieved for the CaMn (**a, b**) and HET-s (**c, d**) motifs in the cross-validation experiment. Performance on training and validation sets without smoothing (**a, c**) and on validation set with smoothing (**b, d**) is shown. Data for evolutionary learning without (with) contact constraints are plotted in shades of orange (green). Data for statistical learning without (with) contact constraints are plotted in shades of blue (magenta). Note logarithmic scale for duration of training.

Analogous tests were performed on the larger and more diverse HET-s set. While GA again needed on average 3000-4000 steps to converge, IO needed 200-300 iterations with the convergence criterion calculated over 100 iterations (Fig. 1c). The statistical learning led to the maximum average performance of 0.79, in comparison to 0.63 achieved with the evolutionary scheme. The contact constraints still sped up the convergence of IO, but at the same time they limited the top performance on the training *and* validation sets by around 0.15. This was in contrast to GA-trained grammars which benefited from the contact constraints by around 0.10. The over-fitting gap was moderate: around 0.10 for IO and 0.06 for GA. Consequently, the effect of smoothing was very limited (0.01-0.02, Fig. 1d shows results for the PAM10 smoothing).

#### Grammar size

Saturation of performance on the HET-s training set together with the lack of significant over-fitting suggested that grammar size was inadequately small with regard to the diversity of the sample. This prompted us to investigate using larger numbers of non-terminals with the IO algorithm. We opted for relatively long evolution counting on the smoothing to fight back the negative effects of over-fitting. Specifically, we trained rule probabilities of grammars in the CFC form [36, 32] made from 5, 7 and 10 lexical and 10 to 33 structural non-terminals. Consequently, the smallest covering grammar *l*5*s*5 counted 1225 rules, while the largest *l*10*s*30 had 138 200 rules. The stop condition was set to 1.0005 over 100 iterations. At the end of training, grammars retained on average only 198 to 650 rules (117 to 482 with probabilities over 1*e* − 5) including 41 to 151 lexical rules (27 to 60 with probabilities over 1*e* − 5), see Fig. 2b. Of note, the trained grammars with more structural variables had less lexical variables, which appeared to be decreasing asymptotically to 20 (or one lexical rule per one amino acid). Only grammars with least variables did not achieve virtually perfect fit to the training sample (mean AP over 0.999). The performance over the validation set without smoothing dropped for grammars with a medium number of non-terminals, presumably due to the over-fitting (Fig. 2a). Interestingly, the validation performance improved again for grammars with large number of non-terminals. The most plausible explanation is that due to more laborious convergence, the training stopped before significant over-fitting took place. The smoothing of lexical rules probabilities using the PAM 10 model led to the mean AP over the validation set around 0.99 for all grammars with 10 (15) or more structural variables trained without (with) contacts (Fig. 2a).

**Figure 2:**
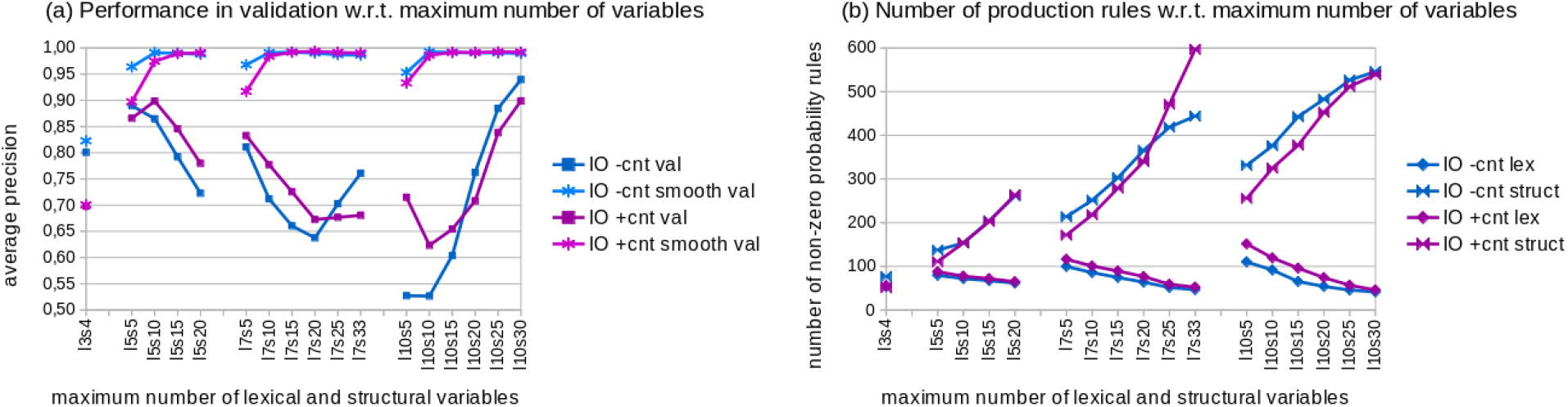
Number of variables effects on performance and grammar size. Mean average precision (**a**) and mean number of lexical and structural rules with non-zero probability (**b**) in the cross-validation experiment are shown with regard to the maximum number of variables (*l*: lexical, *s*: structural). Data for learning without (with) contact constraints are plotted in shades of blue (magenta).

#### Practical performance

It can be reasonably expected that by using grammars learned for the HET-s r1 and r2 motifs, the r0 motifs can be automatically extracted either from random full length sequences (approximated by *PDBfull*) and from NLR proteins (approximated by *NLReff*). Moreover, grammars have to distinguish amyloid signaling motifs from non-amyloidogenic disordered proteins *DisProt*. For the assessment, we used the measures of recall (sensitivity) of the positive sequences at false positive rates (FPR) of 0.01 and 0.001 (the latter for *NLReff* only), and the average precision (AP) against the *DisProt*. To avoid overestimating performance, the minimum observable FPR due to the negative set cardinality was assumed in averaging over the folds (*PDBfull*: 1.2*e*−3, *NLReff* : 3.7*e*−4). The tests were conducted for *l*7*s*15 grammars and neighboring configurations (*l*5*s*15, *l*7*s*10, *l*7*s*15, *l*7*s*20 and *l*10*s*15).

In the scenario with the r0 motifs searched among non-prionic *NLReff*, the mean recall was in range 0.66-0.73 at FPR of 1*e*−3, and in range 0.79-0.84 at FPR of 1*e*−2. Best performance was achieved with *l*5*s*15 grammars trained without the contact constraints. In the search against *PDBfull*, the mean recall at FPR of 1*e*−2 was in range 0.70-0.76. The mean AP against *DisProt* was between 0.96-0.98. Considerable improvement was achieved when scores from grammars obtained in several runs under the same conditions were averaged before classifying hits. The effect was most pronounced were moving from a single grammar to a pair, but increase was notable at least up to 6 grammars combined (the largest number tested was 8). In this case the mean recall was in range 0.82-0.84 at FPR of 1*e* 3 and 0.85-0.91 at FPR of 1*e*−2 for *NLReff*. Similarly, the mean recall at FPR of 1*e*−2 increased to 0.81-0.85 against the *PDBfull*. The already high performance against non-amyloid disordered proteins was kept.

### Application to bacterial amyloid signaling motifs

Performance of the method was further tested on newly identified bacterial amyloid signaling motifs (*BASS*) [13]. A major difference with regard to the previous tests is the variable length of sequences in the BASS families. The variation could result either from indels, intra-family diversity, or motif truncation in the course of extraction. While sequences in each family were in general alignable, which was exploited in the contact constraints prediction, it is worth noting that the PCFG input was unaligned. We used the set-up established for HET-s and the *l*7*s*15 covering grammar.

#### Cross-validation

We first applied the 6-fold standard cross-validation scheme to compare performance of grammars trained with and without contact constraints. In the validation phase, positive sets (variable length) and the negative set *PDBfrag* (made of 40-amino acid chunks) were scanned with the grammars using the 20-to-40-amino-acid window. Since the average precision is sensitive to the cardinality ratio of positive and negative sets, we used the Youden’s Index [78], as a complementary measure, comparable between various BASS families. The results are shown in columns AP and YI in Tab. 1.

**Table 1:**
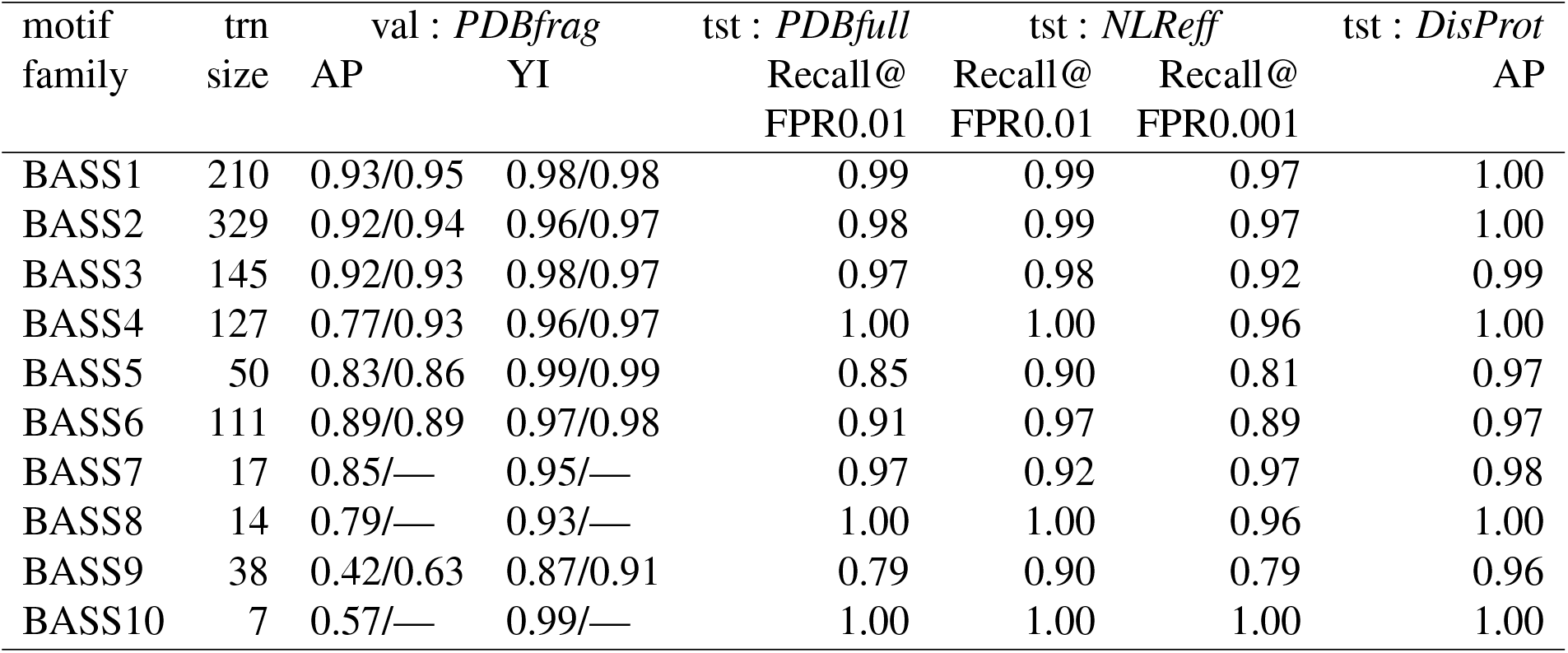
Average performance of grammars for individual ASMs. AP and YI are given for both training without contacts / with contacts. The test set performance is shown only for the mode with the best validation AP. Notations: trn, val, tst are training, validation and testing positive sets, AP is the average precision and YI is the Youden’s Index.

The cross-validation experiment showed that grammars learned the pattern in all cases. The mean Youden’s index ranged from 0.87 to 0.99, which corresponded to the mean average precision from 0.42 (BASS9) to 0.96 (BASS1). While using the contact constraints was favourable in terms of average precision whenever available, the effect was substantial only for BASS4 (increase from 0.77 to 0.93) and BASS9 (increase from 0.42 to 0.63). The best performing constraints comprised of 1 to 3 pairs of residues in contacts.

#### Practical performance

Next, we evaluated performance in the practical settings already defined for the HET-s set. The results are shown the last four columns in Tab. 1. At FPR of 0.01, the grammars found at least 90% of NLR-side motifs among non-prionic NLR effector domains, and at least 79% among sample PDB sequences. At FPR of 0.001 against *NLReff*, the grammars found at least almost 89% of NLR-side motifs for all families except BASS5 and BASS9 (around 80%). Also, grammars allowed for distinguishing ASMs from non-amyloidogenic disordered regions (*DisProt*) with the average precision ranging from 0.96 to 1.00. Overall, single grammars for individual BASS families typically performed better then combined grammars (using the score averaging scheme) for the HET-s motif.

### Generalization of bacterial amyloid signaling motifs

Having checked that the PCFG framework can effectively model each motif family, we aimed at assessing whether the method can be used to obtain a general model of the amyloid signaling motifs.

#### Bacterial motifs

The combined set of all ten BASS families was used to train universal BASS grammars within the 6-fold cross-validation scheme (the folds were made by merging the folds for each motif family). The number of symbols used in the grammars was 7 or 10 for lexical variables and 15, 20 or 30 for structural variables, as it can be reasonably expected that a grammar covering several families requires more complex structures.

The performance of resulting grammars in cross-validation ranged from AP of 0.79 for *l*7*s*15 trained without contact constraints to 0.86 for *l*7*s*30 trained with contact constraints. The best performance was achieved at the log score threshold of 2.4 to 2.9 yielding the Youden’s index of 0.87 to 0.92. In the cross-validation experiment, adding more lexical and structural symbols, and using the contact constraints improved AP of grammars. The same held for practical performance evaluation on BASS-containing versus non-prionic N-termini of NLRs, except that the benefit of training with the constraints was weaker (Fig. 3a). The best grammars accepted up to around 85% (94%) of the positive test sample at the false positive rate of 1*e*−3 (1*e*−2) against *NLReff*. The mean average precision against the non-amyloidogenic disordered proteins was in the range of 0.988-0.995. We did not notice significant trends with regard to the stop condition.

**Figure 3:**
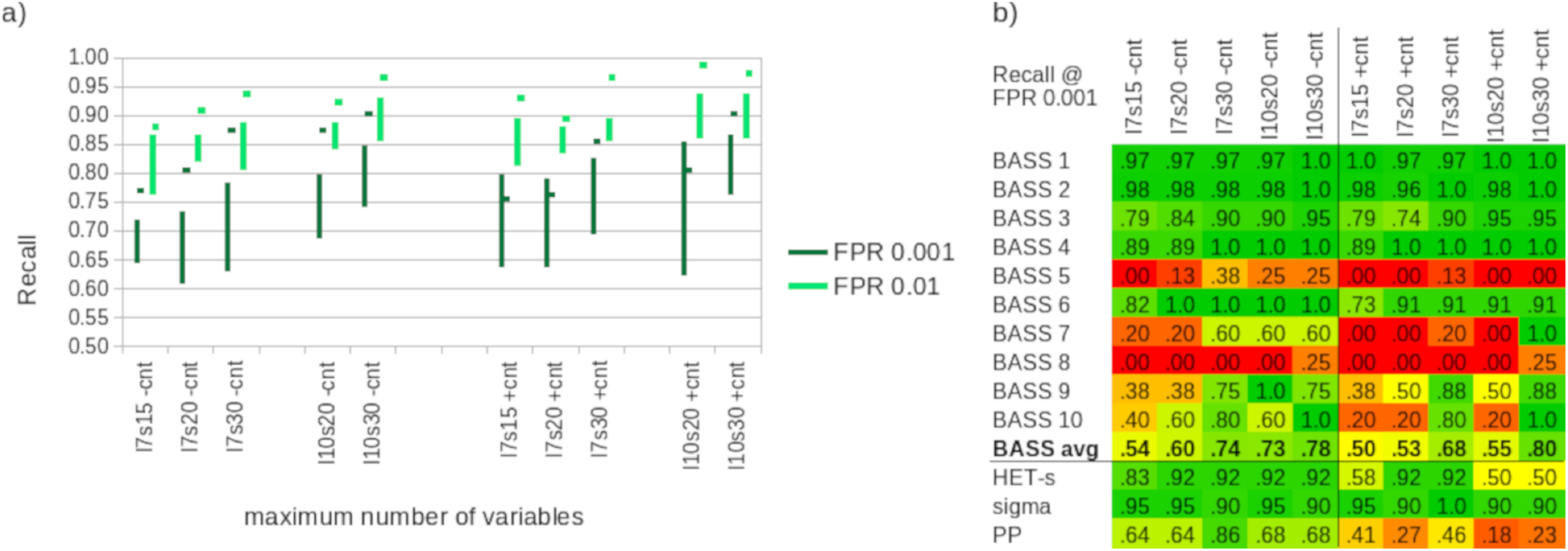
Performance of the generalized BASS grammars on bacterial and fungal test motifs from NLR proteins. (**a**) Recall at the false positive rates of 1*e*−2 (pale green) and 1*e*−3 (dark green) for BASS 1-10 motifs with regard to the maximum number of variables in grammars (*l*: lexical, *s*: structural) and presence/absence of the contact constraints (±*cnt*). Vertical bars indicate recall ranges achieved using single grammars in six runs. Short horizontal bars indicate recall obtained using the score averaging scheme. (**b**) Recall at the false positive rates of 1*e*−3 obtained using the score averaging scheme for each BASS class separately and for the three fungal amyloid signaling motifs.

#### Fungal motifs

Encouraged with the results for the test BASS set, we were curious if the grammars were general enough for searching for novel motifs. Thus, we tested the all-BASS grammars on fungal NLR N-termini with HET-s, sigma and PP/ses_B motif instances against the non-prionic NLR N-termini. The experiment showed considerable performance. The sigma motif was readily distinguishable with the BASS grammars (average recall of 67-87% at FPR of 1*e*−3), followed by the HET-s motif (recall 25-63%), and the PP motif (recall 11-46%). Very good performance with the sigma motifs was likely due to the relatively high length and amino-acid composition similar to some bacterial motifs (e.g. BASS 2, 3 and 7). On the other hand, quite fair sensitivity with the quite distinctive HET-s, as well as relatively poor sensitivity to the RHIM-like PP (seemingly related to BASS3), cannot be easily explained. Unlike the previous BASS tests, training with constraints tended to lower performance for the fungal ASMs, as did adding more symbols and longer training. (With the exception of the sigma motif, with universally high performance.) This is not unexpected, since extrapolating outside the training domain has to carefully avoid over-fitting to be successful.

#### Score averaging

We also noticed high variation of performance of grammars trained with the same parameters on different folds. While recall (at FPR of 1*e*−3) on the NLR-side BASSes in most cases varied only by 10-15% (Fig. 3a), it differed from zero to 75% on the HET-s motif and from zero to 46% on the PP motif test set. While this could be partially due to smaller test sets, it also clearly suggested sub-optimal character of individual grammatical models. Thus, we resorted to the strategy of averaging scores from several grammatical classifiers when scanning sequences. Following previous experiments with grammars for individual motifs, we used average scores of six grammars (one from each training fold). The procedure yielded very good results, with the recall at FPR of 1*e*−3 increasing up to 86-92% for all fungal amyloid signaling motifs for the *l*7*s*30 setup trained without contact constraints (Fig. 3b). Performance of grammars trained with the contact constraints was still rather poor for PP and somehow mixed for HET-s, in contrast to universally good performance of grammars trained without the constraints.

The averaging approach also improved the performance on NLR-side BASSes, up to recall of around 90% (95%) at FPR of 1*e*−3 (1*e*−2), as indicated with short horizontal bars on Fig. 3a. In fact, the averaging scheme most often outperformed the best single grammars. The breakdown of the results by BASS class indicates that the generalized BASS grammars fairly well recognized BASS 1-4 and BASS 6 test samples (recall of 0.90 at FPR of 1*e*−3 for averaging over *l*7*s*30 grammars or larger). Performance over classes BASS 7, 9 and 10 was mixed as grammars with 30 structural variables were doing much better than smaller ones. BASS 5 and BASS 8 classes were not modeled properly (Fig. 3b).

On the set of other BASS motifs, the averaging approach resulted in recall from 50% to 75% (73% to 89%) at FPR of 1*e*−3 (1*e*−2) against *NLReff*. As with the fungal motifs, grammars trained without the contact constraints performed better.

#### Pairing potential

In order to assess the structure of grammatical descriptors of fungal motifs, we scanned the C-terminal 80 amino-acid fragment of HET effector sequence (accession: Q03689) and the N-terminal 50 amino-acid fragment of its genomic neighbor NLR sequence (accession: CAL30199) using the score averaging approach and the 20-to-40 amino-acids window (Fig. 4a). In CAL30199, the best matches very well covered the r0 motif. In the case of Q03689, the best matches were centered on the loop between the r1 and r2 motifs. Grammars trained without the contact constraints partially overlapped the motifs, while grammar trained with the contact constraints covered the r1 motif using the maximum window size of 40 (except *l*10*s*30). A plausible explanation of discrepancy with the actual motif positions is high content of charged residues in mutually complement r1 and r2 motifs, which is atypical for the BASS motifs. Then, we compared the probability maps of parsing of the best matching sequence fragments with their corresponding spatial distance maps. For the NLR side CAL30199, we used the r2 motif fold, as often assumed in literature [54] (Fig. 4b). The pairing maps generated with grammars trained without the contact constraints were dominated by an apparently artifactual antidiagonal signal resembling the pattern observed previously in mostly likely parse trees ([32]). The signal was partially present also on pairing maps generated with grammars trained with the contact constraints. However, in this case, there was also a clear signal corresponding to actual structures of the HET-s fold: the "bulge" from A228 to I231, the tip of the main loop from T233 to V239, and the second loop from Q240 to V244 (positions according to the r1 motif, see Fig. 1 in [54] for reference).

**Figure 4:**
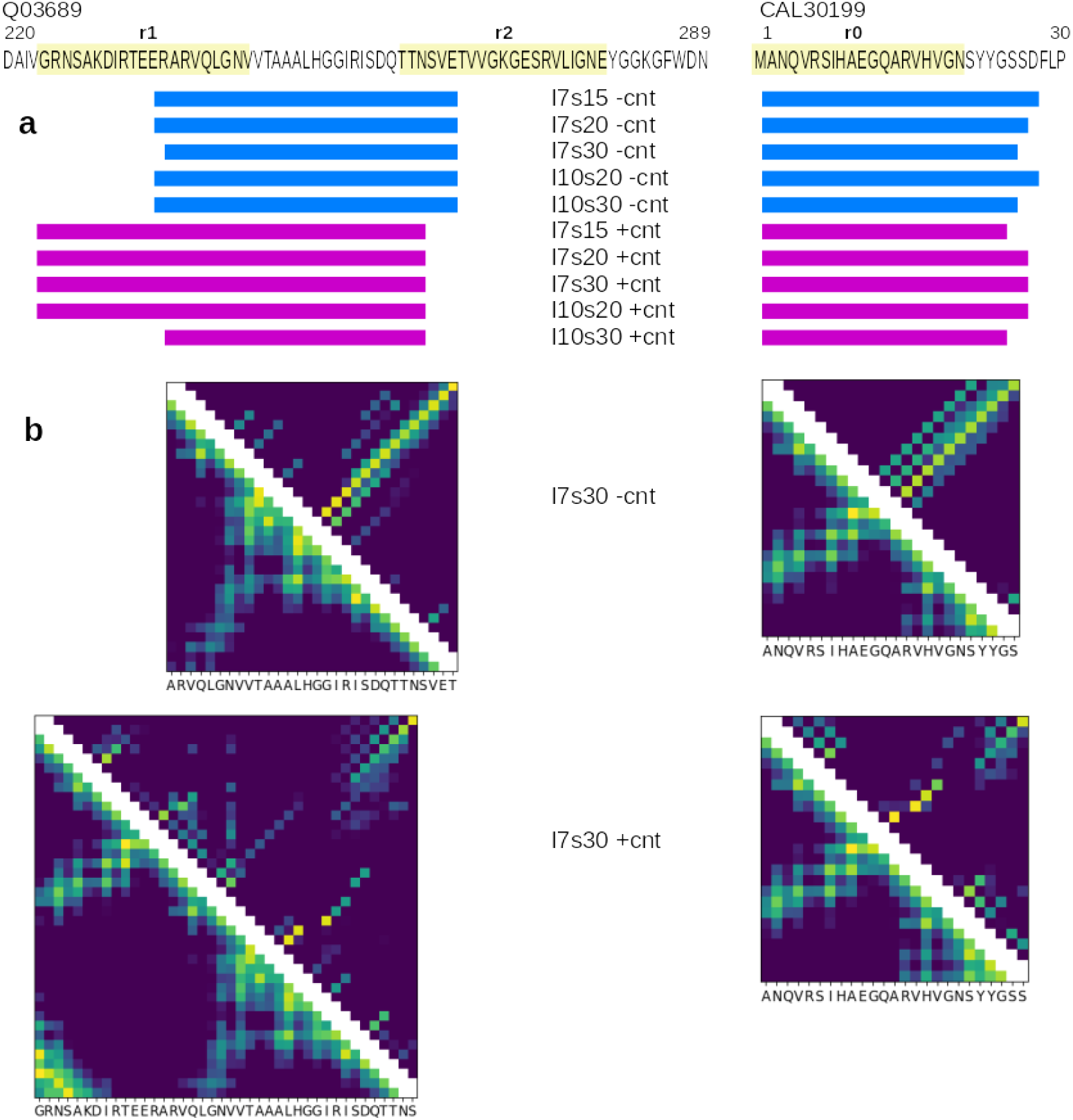
Qualitative analysis of generalized BASS grammars match to experimentally solved HET-s structure pdb:2kj3 [56]. (**a**) Best matches in effector-side C-termini and NLR-side N-termini sequences. Actual position of r1, r2 and r0 motifs are marked in pale yellow. (**b**) Comparison of the structure-derived distance map (lower left triangle, color scale from 4 Å or less (yellow) to 16 Å or more (blue) according to the C*β* distance) to grammar-derived pairing probability map (upper right triangle, logarithmic color scale from 0.01 or less (blue) to 1 (yellow)).

#### Lexical rules

Grammars are considered as human readable descriptors. We analyzed grouping of amino acids according to high probability of being rewritten from given lexical non-terminal symbols (above 0.1). We focused on groupings preserved in at least half of grammars, for each grammar size and the contact constraints option. Almost universally preserved was single non-terminal dedicated overwhelmingly to glycine. Very often grammars consisted of non-terminal symbols dedicated to alanine (sometimes together with serine) and variables likely rewritten to valine and isoleucine (sometimes together with leucine). Next common association was a non-terminal with emissions dominated by glutamine, an amino acid relatively frequent in the prions. On the other hand, no clear pattern was observed with aspargine and aspartic acid, even though they are seemingly relevant for the amyloid signaling motifs. Many grammars included also a non-terminal symbol likely rewritten to a mix of arginine, glutamic acid, lysine, proline, serine, threonine and in some cases histidine, though partition varied. This subset may correspond to a group present in some classical 5-letter alphabets [79, 80] but without amino acids characteristic to the prions. The groupings defined by PCFGs were also partially similar to the best-performing reduced alphabet for amyloid hexapeptides search from [20], which included groups for glycine alone, isoleucine-leucine-valine hydrophobics, and the lysine-proline-argine mix.

#### Structural rules

For each sequence, usage of every rule can be recorded, which is the amount of probability mass carried through the rule (as calculated for the Inside-Outside procedure) relative to the overall probability of the sequence given the grammar. If this quantity is summed up for each left-hand-side non-terminal, one obtains usage of each non-terminal symbol. Note that the usage of the start non-terminal is always at least one, and possibly higher if the start symbol is used again in some derivations. We calculated usage of structural non-terminals in the best matching sequence fragments that achieved positive log probability ratio for the positive test samples (NLR-side BASSes and fungal motifs) and *DisProt*), averaged over the cross-validation folds. For each test set, we identified non-terminal symbols with usage of at least 1. Then, we compared the test sets in terms of the highly used symbols using the Jaccard distance metric. Since the structural non-terminals represent higher level structures in the grammar (as *noun phrase* in English), similarities in their usage may reflect similarities between motifs. Varying maximum number of non-terminals and the contact constraints option resulted in different sharing of the highly used symbols. As could be expected, smaller grammars resulted in higher shares (50-80%) than large grammars (30-60%). Nevertheless, some clear patterns emerged. BASS 1 mostly shared highly used structural non-terminals with BASS 9-10 and fungal HET-s, and least with *DisProt*. BASS 2 mostly shared its highly used structural non-terminals with BASS 6 and fungal sigma and PP, while least, again, with *DisProt*. Unsurprisingly, BASS 3 mostly matched the PP motif, followed by other fungal motifs. For BASS 4 the closest match was HET-s, while for BASS 5 it was *DisProt*. Interestingly, the BASS 4 motif is relatively often found repeated in the Bell side, as is the case of HET-s. BASS 7 motif mostly shared highly used symbols with PP and HET-s, and BASS 8 — with PP, *DisProt*, sigma, BASS 5 and BASS 7.

#### Comparison to alternative approaches

Next, we tried identifying sequences with BASS-like motifs using several existing tools. It has to be noted that most of them was not designed specifically for searching short amyloid signaling motifs.

- **PrionW** [81] is a web server based on the pWaltz method for detecting prions, which assumes they consists of an amyloidogenic core inside a disordered region, rich in aspargine and glutamine [15]. With default parameters, the tool did not found any prion motif in the BASS and fungal ASM positive test sets. Lowering the pWaltz cutoff to 0.50 resulted in finding 3 out of 54 fungal motifs but also 5 hits in the non-prionic NLR N-termini negative set. Clearly the tool is not sensitive to the BASS-like motifs.
- **ArchCandy** [24] is a method for detecting beta-arches. With the recommended score threshold of 0.56, it marked as positive 50% of sequences in the BASS test set, as well as 83% sequences with the fungal ASM motifs. However, it also detected beta-arches in 61% of the negative non-prionic NLR N-termini set. This is rather not surprising as beta-arches are a broad category comprising also sequences that are neither prionic nor amyloidogenic.
- **PASTA2** [19] is a popular tool for detecting amyloid structural aggregation. With the peptide mode settings, PASTA2 reported amyloid-like aggregation regions in 38% of sequences in the BASS test set and in 67% sequences with the fungal ASM motifs. Interestingly, it also founds such regions in 91% of the negative non-prionic NLR N-termini set. Indeed, it is known that short potentially amyloidogenic regions can be found in proteins never observed to form amyloids [82].
- It is conceivable that combining **ArchCandy** and **PASTA2** may improve the accuracy of BASS search. For each sequence, we checked if any top 20 amyloidogenic region predicted with PASTA overlapped with any beta-arch predicted with ArchCandy, using the default detection thresholds of both tools. The recall was 24% for the BASS test set and 41% for *NLReff*, clearly showing that the approach is not viable for searching BASSes.
- **ClustalOmega** [57] and **HMMER3** [83] together allows for creating a profile HMM for the set of all training BASSes. Surely, making a single multiple sequence alignment for a collection of likely unrelated motifs is at least debatable in terms of good practices in bioinformatics. Nevertheless, we used the *–auto* option of ClustalOmega and obtained an alignment with the length of 184 amino acids, further fed to HMMER3 (with default options) to create a profile HMM (length: 98 amino acids, effective number of sequences: 14.5). Then, we used the profile HMM to find the best domain matches in each sequence. Interestingly, the approach resulted in reasonable recall of 37% (65%) of NLR-side test BASSes at FPR of 1*e*−3 (1*e*−2) against *NLReff*. For the fungal motifs, the recall at FPR of 1*e*−3 was 22% for sigma, and none for the other two. Finally, we tried the averaging approach with ClustalOmega and HMMER3. Multiple sequence alignments and corresponding profile HMM were created for each of six training folds separately. Scores from all profile HMMs were averaged for each sequence (without checking if the best domain matches overlap). The result was surprisingly good — the average profile HMM approach matched the average PCFG method in detecting NLR-side test BASSes (recall of 91% and 94% at FPR of 1*e*−3 and 1*e*−2, respectively). The outcome for the *BASSother* set was slightly worse, with only 57% (73%) positive samples detectable at FPR of 1*e*−3 (1*e*−2). For the fungal motifs, the averaged profile HMM approach offered the perfect recall in the case of sigma, but only 17% and 27% sensitivity in the case of HET-s and PP motif, respectively (at FPR of 1*e*3). A the first glance, this overall good performance of the method seems puzzling, yet it can be regarded as another example of combining suboptimal results into a working solution.

Despite that pWaltz was developed using the HET-s experimental structure [15], it was unable to detect more than a few BASS-like and NLR-related ASMs even with parameter tweaking. ArchCandy and PASTA2 may be good in identifying regions of interest or discriminating beta-arches and amyloidogenic regions, respectively, but, by design, they are not specific enough for motif searches in large data sets. Although the proposed use of the multiple sequence alignment and profile HMMs was evidently abusive with regard to their theoretical background, the approach performed remarkably well within the modeled meta-class. Extrapolating beyond that meta-class, its sensitivity deteriorated significantly to the level of the least accurate PCFGs for *BASSother* and PP or clearly below for HET-s.

### Experimental verification of selected peptides

Eventually, we tested if selected peptides of those identified as ASMs with grammars and other existing methods (Tab. 2), form spatial structures consistent with the known HET-s structure [56] and consistent with the experimentally demonstrated amyloid-like aggregation [8, 84, 85, 86, 13]. First, to tweak and test the experimental setup, we used two *bona fide* effector-side ASM peptides:

**Table 2:** Top: List of peptides selected for experimental verification. Bottom: results of computational methods. For PCFG and pHMM (score averaging) we provide FPR against *NLReff* at which at least a part of the peptide is detected; for PASTA2 — Pasta Energy Units; for AmyloGram and ArchCandy — score ranged from 0 to 1. Arrow indicates if higher or lower value is more positive. Values exceeding the default or suggested thresholds are shown in bold (the thresholds are 1*e* 2 for PCFG and pHMM, −5 PEU for PASTA2, 0.5 for AmyloGram, and 0.56 for ArchCandy). For PCFG, we provide the range of values for grammars of different maximum number of symbols.

| Peptide | Sequence |  |  |  |  |
| --- | --- | --- | --- | --- | --- |
| ORT49035.1_103_123 | VDLRDAKGVQVG DGNVQINRF |  |  |  |  |
| AAS80314.1_349_385 | SFNNLGSGDQFNTPGGTQNINKGGNEVSGGNFYGSVQF |  |  |  |  |
| AEB69175.1_5_29 | KSRFDQRGQKVIGQQIN VAGDATLP |  |  |  |  |
| RDW70414.1_382_421 | GAPANNTSNSIQHNNSGSGHQNSGSGQQNIGTNTGSGQQ |  |  |  |  |
| Peptide | PCFG<br>FPR↓ | pHMM<br>FPR↓ | PASTA2<br>[PEU]↓ | AmyloGram<br>[0-1]↑ | ArchCandy<br>[0-1]↑ |
| ORT49035.1_103_123 | ≤ **0.0004** | **0.009** | -2.5 | **0.63** | 0.379 |
| AAS80314.1_349_385 | < **0.0004** | **0.006** | -1.6 | **0.78** | 0.555 |
| AEB69175.1_5_29 | **0.007**–0.07 | 0.013 | -3.7 | 0.43 | 0.488 |
| RDW70414.1_382_421 | < **0.0004** | 0.017 | -0.9 | 0.26 | **0.599** |

BASS3 RHIM-like motif from *Frankia sp.* ORT49035.1 (positions 103 to 123) [63, 13] and *Nectria haematococca* sigma motif from AAS80314.1 (349 to 385) [64, 12]. Both could be computationally identified using PCFG (at FPR below 1*e* 3) and profile HMM (at FPR below 1*e* 2) — with the score averaging — as well as with AmyloGram [20]. The latter peptide only marginally missed the ArchCandy threshold. Then we analyzed *Methanothrix soehngenii* AEB69175.1 (5 to 29) [65]. This NLR-side motif resembling BASS3 was originally identified through the local pairwise alignment of proteins encoded by neighboring genes, while being missed with the BASS 1-10 HMM profiles search [13]. Here, it could be identified at FPR of around 1*e* 2 with the profile HMM and some PCFG searches; it also scored quite low energy in PASTA2 (yet still above the 95% specificity threshold recommended for peptides). Finally, we experimentally verified a sequence fragment resembling the sigma motif in *Coleophoma crateriformis* NLR protein RDW70414.1 (382 to 421) [66]. The motif is positive according to ArchCandy and PCFGs (FPR below 1*e*−3), while clearly negative according to PASTA2 and AmyloGram ((Tab. 2).

ATR-FTIR spectra of the four selected peptides (see Materials) in dried form and D_2_O solutions with corresponding second derivative spectra in amide bands range (1750-1500 cm^−1^) are presented in Fig. 5a. Spectral characteristic for peptides ORT49035_103_123, AAS80314_349_385 and AEB69175_5_29 are typical for aggregates. The Amide I’ band maxima are located below 1635 cm^−1^, what is characteristic for the *β*-cross structures. While, in ATR-FTIR spectrum of peptide RDW70414_382_421 no spectral signatures of aggregation process were found. More accurate information about studied peptides can be obtained from the second derivative and decomposition of the Amide I’ band into sub-bands (Fig. 5b). These processes clearly revealed the complex structure of peptide ORT49035_103_123. The Amide I’ band can be separated into five components: 1702, 1672, 1652, 1631 and 1617 cm^−1^, which can be assigned in order to *β*-sheet or turn, *β*-turn, *α*-helix or extended random or loops, *β* - sheet and aggregates [87, 67]. The origin of component 1652 cm^−1^ is not clear, because the loop and the helical absorption bands overlap in this region [88]. For the other peptides less components in the Amide I’ range were observed, but their assignment is similar.

**Figure 5:**
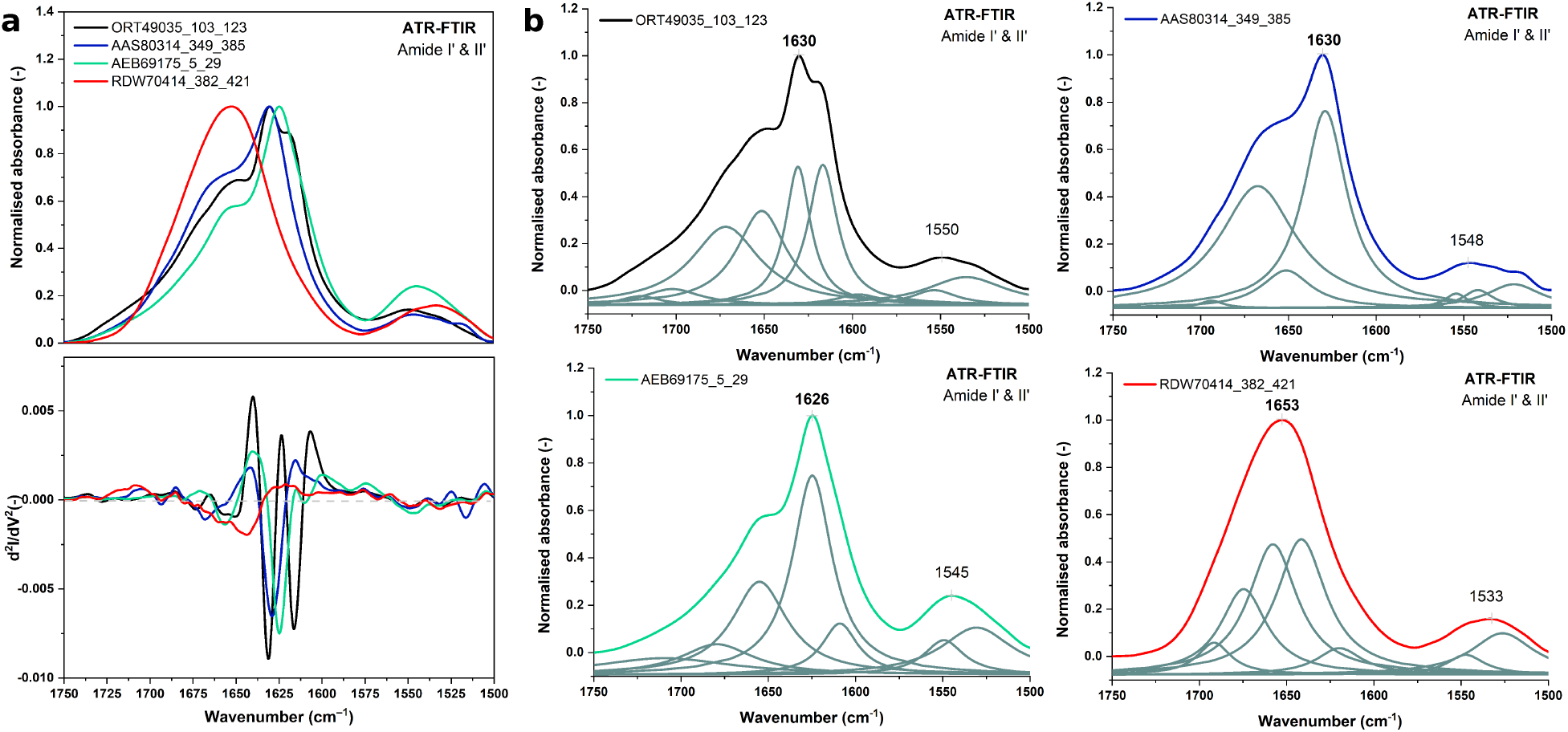
Normalized ATR-FTIR spectra of air-dried peptides films with second derivative spectra in the amide bands region (1750 - 1500 cm^−1^) (**a**) and particular ATR-FTIR spectra with sub-bands obtained from the curve fitting procedure (Amide I’ and II’ regions) (**b**): ORT49035_103_123 — black line, AAS80314_349_385 — blue line, AEB69175_5_29 — green line, and peptide RDW70414_382_421 — red line.

Taken together, the experimental results are compatible with presence of the beta-arch structure and amyloid-like aggregation of peptides ORT49035_103_123, AAS80314_349_385 and AEB69175_5_29. This supports the hypothesis that bacterial, archaeal and fungal NLR-related ASMs share similar structural features. On the other hand, no sign of the amyloid like aggregation was observed for RDW70414_382_421. However, since decomposition of the Amide I’ band for monomeric RDW70414_382_421 reveals similar components to other three peptides, it is likely that it assumes the beta-arch structure as well. Thus, this peptide apparently represents a false positive hit of the computational method, presumably due to over-generalization.

## Discussion

In this piece of research, we addressed some of the previously identified challenges in inferring probabilistic context-free grammars for protein motifs [32]. Overall, presented results show a clear advantage of the Inside-Outside training procedure followed with the lexical probabilities smoothing over the previously used evolutionary scheme [29, 32] in learning of the probabilistic context-free grammars for protein motifs. Current procedure allows for generating much larger grammars with sufficient numbers of non-terminal symbols (e.g. 40). Importantly, the inference time with the IO training is relatively short, ranging from a couple of minutes to a couple of hours, depending on the covering grammar size (on 12 cores of the Intel Xeon E5 (Haswell) machines). In accordance with the literature [34], the obtained grammatical models are typically not globally optimal. In practice, we addressed this by combining (averaging) scores of several individual grammars. The averaging scheme turned out to be very effective, usually outperforming best individual grammars in homology searches (Fig. 3a). Yet, it remains a goal for the future research to optimize learning, e.g. by enhancing the statistical learning with the contrastive estimation [89] whenever possible and combining with heuristic approaches [44, 90].

Recently, we introduced the use of the contact constraints based on known or predicted spatial proximity of residues to learning PCFGs [32]. Due to the context-freeness of grammars and simplicity of the contact rules in the Chomsky Form with Contacts, only a subset of residue-residue contacts can be used in the training. The contacts are properly chosen if there exist correlations between involved amino acid species that are relevant to modeled structures and functions. In such a case the contact constraints facilitate learning through confining the search space towards the most capable solutions. Obviously, the constraints reduce the amount of information that grammars can learn: correlations incompatible with the constraints cannot be captured in the grammar. So, if the constraints are not properly chosen, they may effectively lead to less capable models. This is not so much a problem when modeling samples of highly homologous sequences where the common signal shared by all sequences is very strong. However, in the case of generalizing over multiple motif families, a suboptimal choice of the constraints is likely to hamper the quality of the model more harshly. This could contribute to weaker performance of the generalizing grammars trained with the contact constraints when tested on *BASSother* and some fungal motifs samples (Fig. 3b). Nevertheless, using the contact constraints for training apparently improved compatibility of the residue pairings generated with grammar using the contact rules with the actual protein distance map for the only experimentally solved structure of signaling amyloid, the HET-s motif (Fig. 4b).

One obvious limitation of the PCFG approach is context-freeness: the property that allows a grammar for considering in a single derivation only non-overlapping nested and branching dependencies. Yet, the probabilistic parsing of a sequence consists on scoring over all possible derivations, therefore it is capable of covering several overlapping sets of nested (anti-parallel) and branching dependencies. The PCFG model is, however, not suitable for capturing crossing (parallel) dependencies. This is a serious limitation in the context of modeling protein sequences. Unfortunately, more expressive grammar formalisms are also more computationally expensive. For example, computational time complexity for parsing of the mildly context-sensitive linear indexed grammars [91, 92], which can capture some crossing structures, is 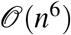. Some alternative are methods building models from the multiple sequence alignment, such as undirected graphical models or Potts models, which can capture information conveyed in the crossing inter-position correlations [93, 94]. There is ongoing research on using such models for aligning sequences in homology searches while avoiding the combinatory explosion [95, 96, 97, 98, 99]. Interestingly, one Potts-based tool under development has been hitherto outperformed by a PCFG-based tool in RNA homology searches [99]. Even if further development leads to unleashing the power of Potts-based models, the requirement of MSA for inferring their parameters makes them rather unsuitable for modeling meta-families of motifs whose members do not share relevant homology yet still share structural or functional principles.

## Conclusions

The results obtained in this piece of research show that the proposed method can infer a model capable of generalizing over a diverse set of families of amyloid signaling motifs. While the profile HMMs remain the method of choice for modeling homologous sets of sequences, PCFGs seem more suitable for building meta-family descriptors and extrapolating beyond the seed sample. (Even if the generalization comes at some price as exposed by the experimentally verified false positive hit.) Indeed, with the score averaging scheme, PCFGs trained without contact constraints outperformed profile HMMs when the BASS-trained models were applied to fungal HET-s (sensitivity of 83-92% *vs.* 17% at FPR of 1*e*−3) and PP motifs (sensitivity of 64-86% *vs.* 27% at FPR of 1*e*−3). Without the averaging, single grammars very clearly outperformed single profile HMMs on both bacterial and fungal motifs. In practice, one can expect even higher specificity of both machine learning methods, since in reporting the results we assumed the conservative upper estimate of the false positive rate when all positive samples scored above every negative sample.

## Supporting information

Data archive

Peptide analytical data

## Abbreviations

AP: Average Precision
ATR-FTIR: Attenuated Total Reflection – Fourier Transform In-frared
BASS: Bacterial Amyloid Signaling Sequence
BLOSUM: Blocks Substitution Matrix
CFC: Chomsky Form with Contacts
CNF: Chomsky Normal Form
DIC: Dissolved Inorganic Carbon
DMF: Dimethylformamide
Fmoc: 9-fluorenylmethoxycarbonyl
FPR: False Positive Rate
GA: Genetic Algorithm
HMM: Hidden Markov Model
HPLC: High-Performance Liquid Chromatography
IO: Inside-Outside algorithm
MS: Mass Spectrometer
MSA: Multiple Sequence Alignment
NBS-LRR: Nucleotide-Binding Site – Leucine-Rich Repeats
NLR: an umbrella term for NOD-Like Receptor and NBS-LRR
NMR: Nuclear Magnetic Resonance
NOD: Nucleotide-Oligomerization Domain
PAM: Point Accepted Mutation
PCFG: Probabilistic Context-Free Grammar
PDB: Protein Data Bank
PEU: PASTA Energy Unit
pHMM: profile HMM
ASM: Amyloid Signaling Motif
RNA: Ribonucleic Acid
TFA: Trifluoroacetic Acid
TIS: Triisopropylsilane
TOF: Time Of Flight
YI: Youden’s index

## Declarations

### Ethics approval and consent to participate

Not applicable.

### Consent for publication

Not applicable.

### Availability of data and materials

The source code is available at git.e-science.pl/wdyrka/pcfg-cm under the GNU General Public License v3.0. The datasets used in the current study and the peptide analytical data are included as supplementary information files.

### Author contributions

WD conceived the study, designed, performed and analyzed the computational experiments, prepared figures and tables, authored the manuscript, developed the software. MG-G designed, performed and analyzed the spectroscopy experiments, prepared figures, authored the manuscript. MS designed and performed the peptide synthesis, authored the manuscript. All authors read and approved the final manuscript.

## Acknowledgements

Not applicable.

