## Supplemental Figure 1 for "Searching for universal model of amyloid signaling motifs using probabilistic context-free grammars"

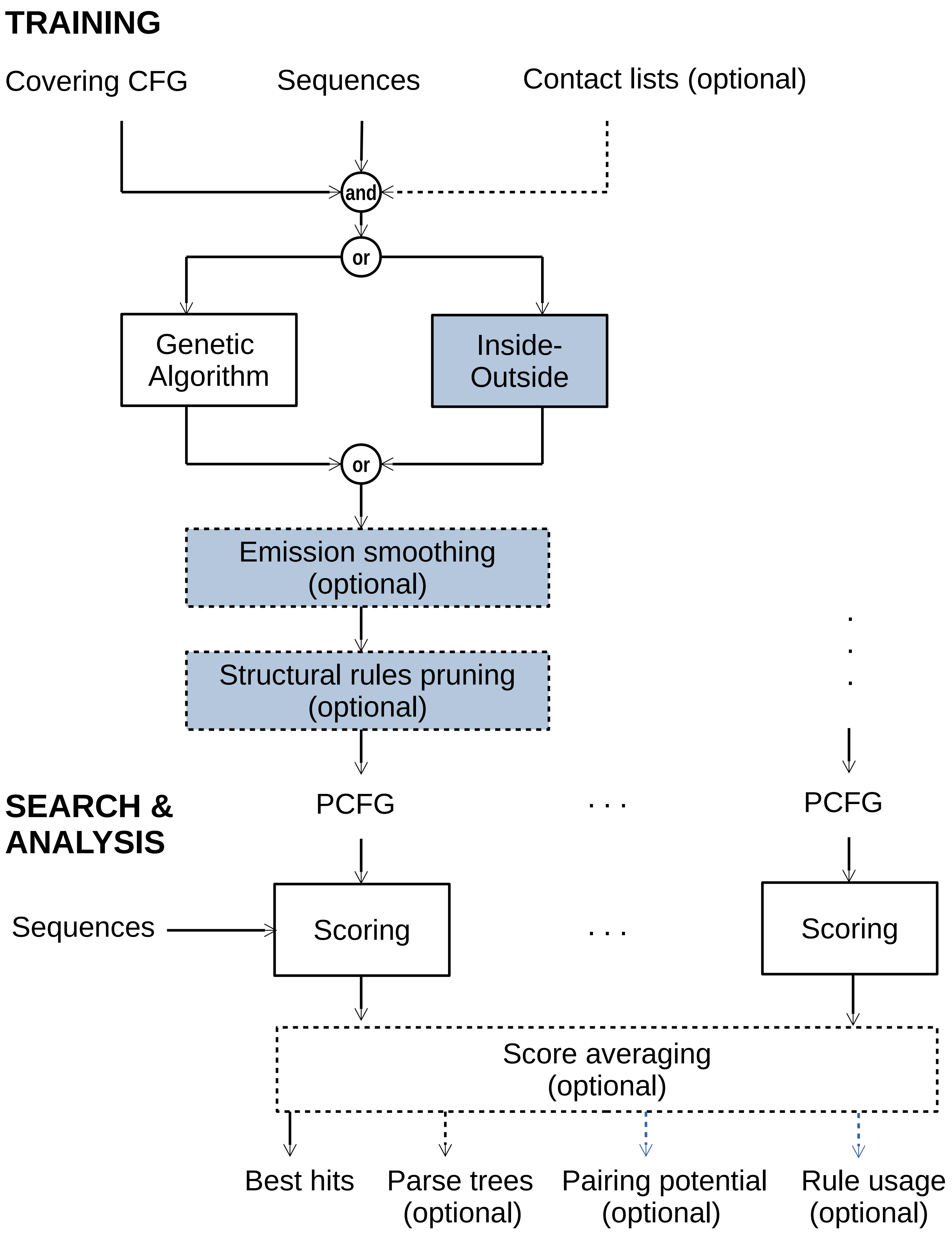


Supplementary Figure 1: The outline of the processing pipeline in the probabilistic context-free grammar-based framework for protein sequence analysis. Optional features, inputs and outputs are marked with dotted lines. Newly introduced features and outputs are marked with grey blue.
