## Supplementary material for "Searching for universal model of amyloid signaling motifs using probabilistic context-free grammars": Data archive: README

Files in these folders use internal names for the BASS classes.
Here is the mapping:
C01 == BASS1
C02 == BASS2
C03 == BASS3
C06 == BASS4
C07 == BASS5
C09 == BASS6
C12 == BASS7
C20 == BASS8
Cya03 == BASS9
Cnlr26 == BASS10
