## Supplementary material for "Searching for universal model of amyloid signaling motifs using probabilistic context-free grammars": Peptide analytical data

| Name | Full name | Formula | $M_{cal.}$[1] | $M_{MS}$[2] | HPLC $t_{ret.}$[3] [min] |
|---|---|---|---|---|---|
| PPT_1 | ORT49035.1_103_123 | $C_{98}H_{163}N_{33}O_{31}$ | 2300.2<br>[1/2M+1] 1150.62<br>[1/3M+1] 767.42<br>[1/4M+1] 575.81 | [1/2M+1] 1150.62<br>[1/3M+1] 767.06<br>[1/4M+1] 576.05 | 8.840 |
| PPT_4 | SesB_349_385 | $C_{168}H_{246}N_{50}O_{59}$ | 3910.83<br>[1/2M+1] 1955.90<br>[1/3M+1] 1304.27<br>[1/4M+1] 978.46 | [1/2M+1] 1955.94<br>[1/3M+1] 1304.27<br>[1/4M+1] 978.43 | 8.350 |
| PPT_5 | AEB69175.1 5 29 | $C_{117}H_{197}N_{39}O_{36}$ | 2726.8<br>[1/2M+1] 1363.75<br>[1/3M+1] 909.50<br>[1/4M+1] 682.38 | [1/2M+1] 1363.75<br>[1/3M+1] 909.50<br>[1/4M+1] 682.38 | 7.763 |
| PPT_6 | RDW70414_382_421 | $C_{146}H_{232}N_{58}O_{63}$ | 3808.6<br>[1/2M+1] 1904.85<br>[1/3M+1] 1270.23<br>[1/4M+1] 952.93 | [1/2M+1] 1904.98<br>[1/3M+1] 1270.24<br>[1/4M+1] 952.93 | 7.841 |

[1] $M_{cal.}$ – calculated mass of the peptide
[2] $M_{MS}$ – found mass of the peptide using HRMS
[3] HPLC $t_{ret.}$ – retention time in analytical HPLC spectra
